# Diet influences early microbiota development in dairy calves without long-term impacts on milk production

**DOI:** 10.1101/408898

**Authors:** Kimberly A. Dill-McFarland, Paul J. Weimer, Jacob D. Breaker, Garret Suen

**Author notes:** Address correspondence to Garret Suen, Department of Bacteriology, University of Wisconsin-Madison, 5159 Microbial Sciences Building, 1550 Linden Drive, Madison, WI, USA 53706-1521. Current address: Department of Microbiology and Immunology, University of British Columbia, Vancouver, British Columbia, Canada (Kimberly A. Dill-McFarland).

## Abstract

Gastrointestinal tract (GIT) microorganisms play important roles in the health of ruminant livestock and impact production of agriculturally relevant products, including milk and meat. Despite this link, interventions to alter the adult microbiota to improve production have proven ineffective as established microbial communities are resilient to change. In contrast, developing communities in young animals may be more easily altered but are less well-studied. Here, we measured the GIT-associated microbiota of 45 Holstein dairy cows from 2 weeks to first lactation using Illumina amplicon sequencing of bacterial (V4 16S), archaeal (V6-8 16S), and fungal (ITS1) communities. Fecal and rumen microbiota were correlated to growth and milk production of animals raised on calf starter grains and/or corn silage to determine if early-life diet has long-term impacts. Significant diet-associated differences in total microbial communities and specific taxa were observed by weaning (8 weeks), but all animals reached an adult-like composition between weaning and 1-year. While some calf diet-driven differences were apparent in the microbiota of adult cows, these dissimilarities did not correlate with animal growth or milk production. This suggests that initial microbial community establishment is impacted by early-life diet, but post-weaning factors have a greater influence on adult communities and production outcomes.

**SIGNIFICANCE:** The gut microbiota is essential to the survival of many organisms, including ruminants that rely on microorganisms for nutrient acquisition from dietary inputs toward the production of products like milk and meat. While alteration of the adult ruminant microbiota to improve production is possible, changes are often unstable and fail to persist. In contrast, the early-life microbiota may be more amenable to sustained modification; however, few studies have determined the impacts of early-life interventions on downstream production. Here, we investigated the impacts of agriculturally relevant calf diets, including calf starter and corn silage, on gut microbial communities, animal growth, and production through the first lactation cycle. Thus, this work serves to further our understanding of early-life microbiota acquisition as well as informs future practices in livestock management.

## INTRODUCTION

The mammalian gastrointestinal tract (GIT) houses a diverse microbial community that can significantly impact both the lifetime and evolutionary success of its host (McFall-Ngai *et al.*, 2013). These microbial communities contribute to host survival by extracting nutritionally (Bergman, 1990) and developmentally relevant (Gensollen *et al.*, 2016) compounds from dietary inputs. Microbial colonization of the GIT begins early in life (Fonty *et al.*, 1987; Klein-Jöbstl *et al.*, 2014) and once established, gut communities are resistant to change outside of extreme or sustained interventions (Weimer *et al.*, 2010; Yanez-Ruiz *et al.*, 2015). Therefore, efforts to alter the GIT microbiota to improve any number of host outcomes may be most effective during early-life before microbial communities have reached an adult-like steady-state. However, factors impacting microbial acquisition and long-term maintenance in the GIT are not well understood.

In mammals, there is a dramatic shift towards an adult-like microbiota during weaning, likely as a result of changes in diet (for a review, see (Lallès, 2012)). This indicates that diet is a strong contributing factor in gut microbiota establishment and could serve as a tool to directionally alter the microbiota in early-life. Microbiota manipulation through diet is of interest in a number of areas, particularly in agriculture where the use of other therapeutics like antibiotics and probiotics are tightly regulated (Bajagai *et al.*, 2016). In the beef and dairy industries, differences in the microbiota have been linked to bovine milk production (Elie Jami *et al.*, 2014; Jewell *et al.*, 2015), weight gain (Carberry *et al.*, 2012), and methane emissions (Wallace *et al.*, 2015). Strong associations occur between these traits and the rumen microbiota because this enlarged foregut compartment houses the microbial community responsible for the majority of dietary digestion and fermentation (Flint *et al.*, 2008). Therefore, it may be possible to utilize diet to manipulate bovine GIT microbial communities to increase animal performance (Malmuthuge *et al.*, 2015).

Although diet has been investigated as a means of altering the adult cow microbiota (for examples (de Menezes *et al.*, 2011; Patra and Yu, 2012)) and production (for examples, (E Jami *et al.*, 2014; Torok *et al.*, 2014)), calf feed has not been as fully assessed. Previous dietary work in dairy calves focused on weight gain, as this is positively associated with increased downstream milk production (Soberon *et al.*, 2012); however, studies investigating the calf GIT microbiota failed to follow calves through to maturity (Vlková *et al.*, 2008; Cannon *et al.*, 2010; Jatkauskas and Vrotniakiene, 2010; Signorini *et al.*, 2012; Malmuthuge *et al.*, 2013). Given that early-life feed has demonstrated long-term effects on the adult microbiota of other ruminants (Yáñez-Ruiz *et al.*, 2010; Abecia *et al.*, 2013, 2014), and that microbial communities in the pre-weaned dairy calf are highly variable (Jami *et al.*, 2013; Dill-McFarland *et al.*, 2017) and amenable to change (Yanez-Ruiz *et al.*, 2015), we posit that early-life dietary interventions in dairy calves can impact microbial colonization with long-term consequences for the adult cow microbiota and production.

To investigate this, we utilized Illumina amplicon sequencing of bacterial (V4 16S), archaeal (V6-8 16S), and fungal (ITS1) communities of dairy cows from 2 weeks after birth to the middle of their first lactation cycle. We compared animals raised on low-fiber, high-protein calf starter grains (previously investigated here (Dill-McFarland *et al.*, 2017)) to those raised on high-fiber corn silage or a mixture of starter grains and corn silage. We then correlated the fecal and rumen microbiota to outcomes including weight gain and milk production to determine the short- and long-term impacts of calf diet.

## RESULTS

### Sample collection and sequencing

In this study, 9 bulls were retained through weaning (8 weeks) and 33 cows through their first lactation cycle (> 2 years). Three cows were removed from the study prior to 2-year sampling due to poor health (cow 5013), reproductive issues (cow 5055), or a broken leg (cow 5025). Feces were sampled from animals from 2 weeks to their first lactation (> 2 years), while rumen contents were sampled from bulls sacrificed at weaning and a subset of cannulated cows (N = 12) at 1 and 2 years. Feces were not sampled from 5 cows at 2 weeks or 8 cows at 8 weeks. Replacement samples were obtained at 3 weeks but were not taken at 9 weeks, because animals were on a different diet by this time (Table S1). All other fecal (N = 183) and all rumen (N = 162) samples were obtained at their target age (Table S2).

Archaeal, bacterial, and fungal amplicons were sequenced for all rumen samples, and archaeal and bacterial amplicons were sequenced for all fecal samples. Fungal PCR of calf feces failed to yield sufficient DNA and therefore, only 1- and 2-year feces were sequenced for this amplicon. After filtering in mothur, 540 000 (mean 1 500 ± 85 SE per sample) high-quality archaeal, 13.9 million (39 000 ± 1 000) high-quality bacterial, and 5.3 million (23 000 ± 985) high-quality fungal sequences were obtained. Sequence coverage was deemed sufficient by a Good’s coverage greater than 96.5% for all bacterial and fungal communities and most archaeal communities. A total of 16 fecal archaeal communities (2-week: 12, 4-week: 3, 8-week: 1) had low Good’s coverage (0 – 94%), even after repeated sequencing yielded a minimum of 30,000 contigs per sample (Table S2).

### Calf diet correlates with microbiota at weaning

Calves increased consumption of supplemental feeds from birth to weaning (8 weeks) following an exponential trend (Figure S1), and there were no differences in the log of supplement consumption between diet groups (*P* = 0.681, repeated measure linear regression). Calf diet did not significantly correlate with the overall structure or composition of any rumen community at weaning (Figure 1, PERMANOVA, Table S3). However, silage-fed calves had lower bacterial diversity with higher archaeal diversity and fungal richness in rumen liquids; bacterial diversity was also significantly higher in silage-fed rumen solids (Figure 2, Figure S2, ANOVA, TukeyHSD, Table S3). Rumen archaeal communities of silage-fed calves had higher inter-animal variation (PERMDISP, TukeyHSD, Table S3) at 8 weeks.

**Figure 1.**
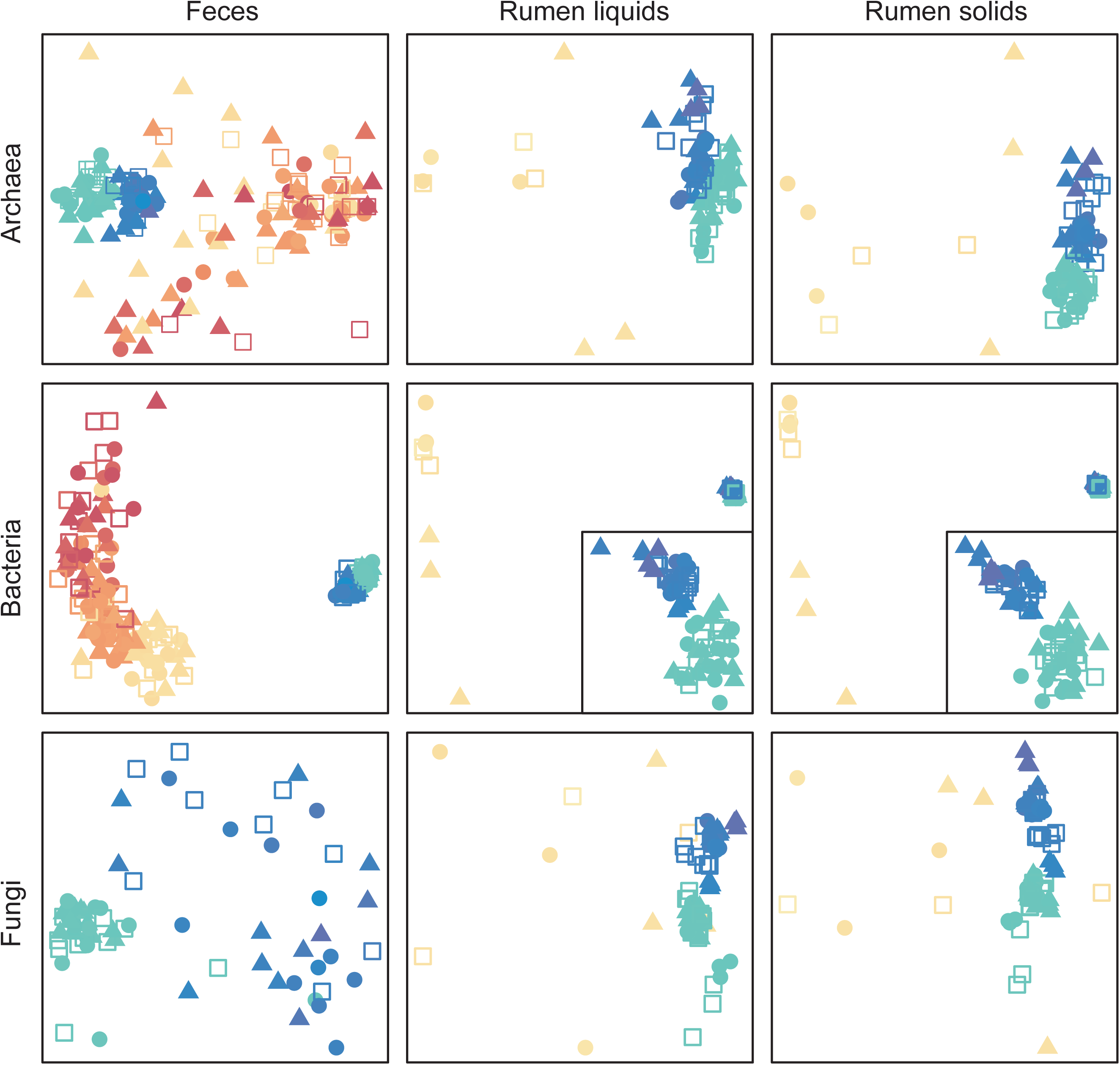
Total fecal and ruminal microbiota in dairy cows raised on different diets. Non-metric multidimensional scaling (nMDS) plots of Bray-Curtis diversity for archaeal, bacterial, and fungal communities in feces as well as rumen liquids and solids are presented. Points are colored by animal age on a continuous rainbow scale at 2-weeks (red), 4-weeks (orange), 8-weeks (yellow), 1-year (green), and 2-years (blue). Calf diet is indicated by circles (calf starter), triangles (corn silage), and open squares (mixed). Bacterial rumen plots have an inset plot displaying the 1- and 2-year clusters in more detail.

**Figure 2.**
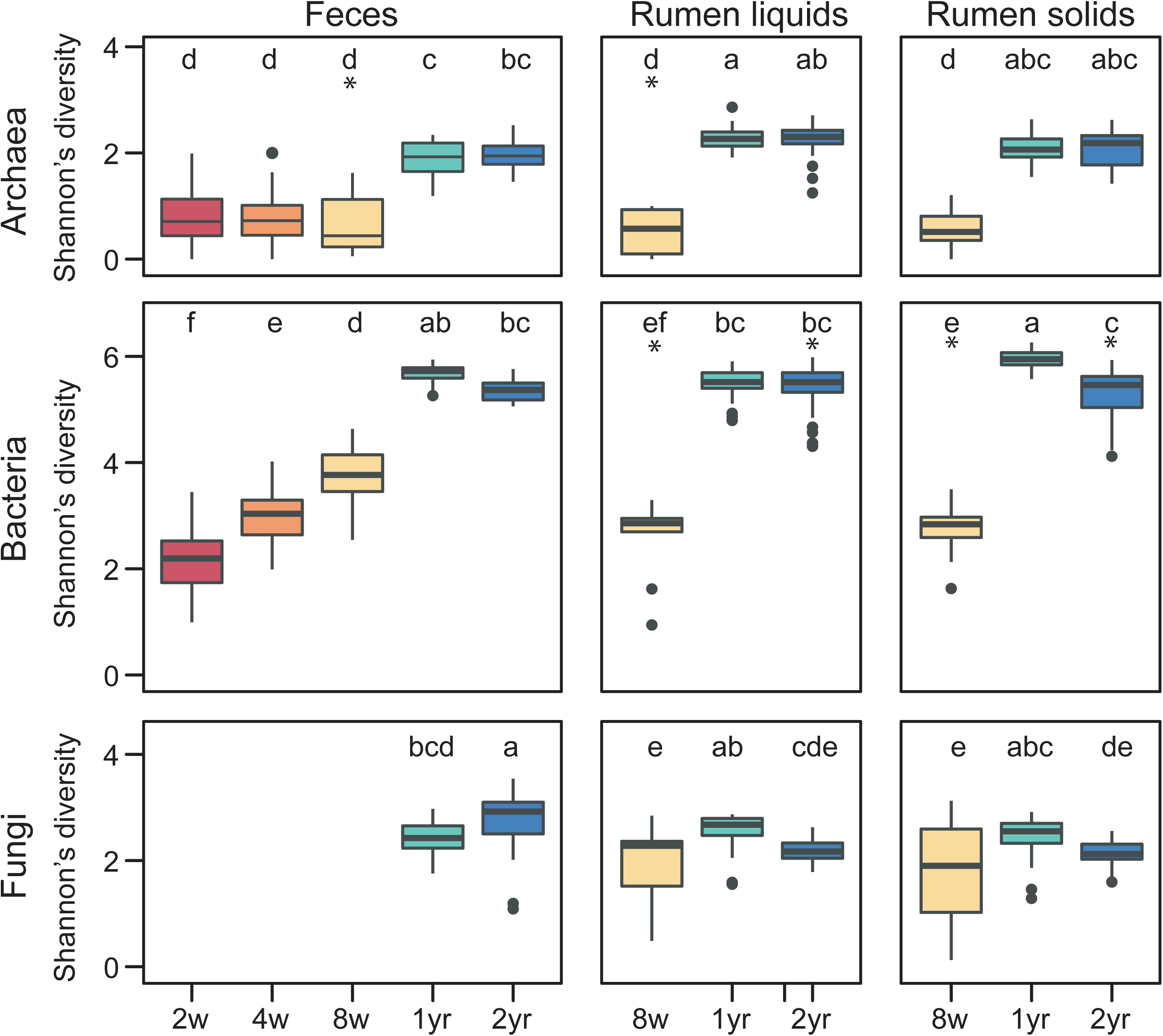
Diversity of fecal and ruminal microbiota in cows raised on different diets. Shannon’s diversity of archaeal, bacterial, and fungal communities in feces as well as rumen liquids and solids are presented. Boxes are colored by animal age at 2-weeks (red), 4-weeks (orange), 8-weeks (yellow), 1-year (green), and 2-years (blue). Ages with significantly different diversity across each amplicon are indicated by different letters. Asterisks denote groups containing significant diet differences (TukeyHSD, *P* < 0.05, Table S3).

Among the three calf diets, the rumen microbiotas of silage-fed calves at weaning had the most OTUs in common with adult communities and starter-fed calves had the least (Figure 3). In particular, the rumen communities of silage-fed animals had higher abundances of a number of adult-associated operational taxonomic units (OTUs) that were absent in starter-fed calves (Table S4). Silage-fed animals also, notably, had lower abundances (0 – 60%) of a calf-associated *Methanobrevibacter* (A-OTU1), which dominated starter and mixed diet animals to greater than 90% but was absent in all adult animals. In contrast, starter-fed and/or mixed diet calves had more highly abundant OTUs at weaning that were not present in adults (Table S4).

**Figure 3.**
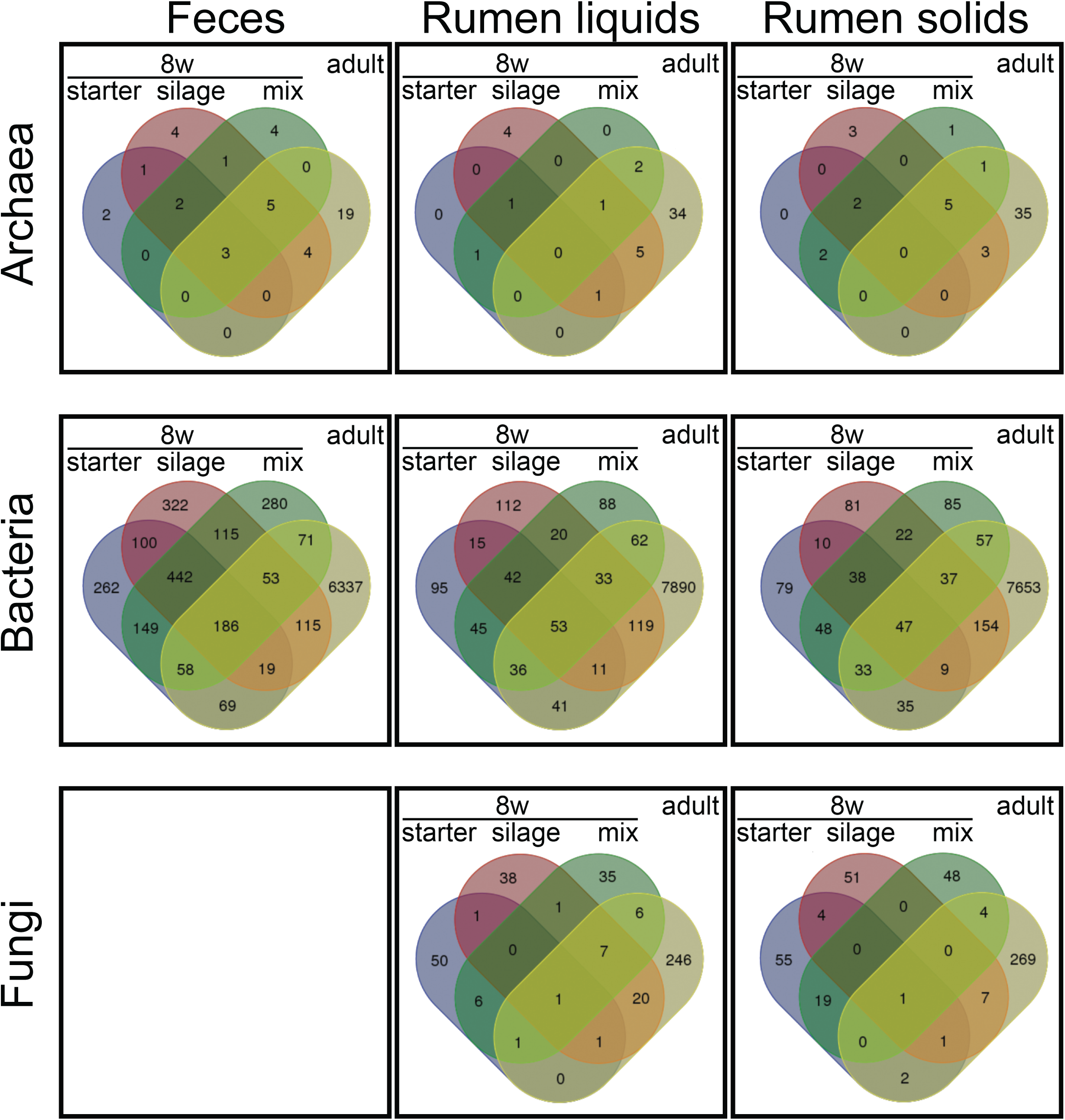
Microbial taxa shared between calves and cows. Venn diagrams of OTUs shared between calves at 8 weeks and adult cows. Calf groups are split by diet, and adults include all samples at 1 and 2 years.

Short chain organic acid (SCOA) concentration profiles in rumen liquids (Table S5) varied with animal age (PERMANOVA, Table S3) and co-varied with the overall archaeal (*P* = 4.95E-02) and bacterial microbiota (Mantel, *P* = 0.010) but not fungi (*P* = 0.129). Calves had significantly higher concentrations and molar fractions of some volatile fatty acids (VFAs), including butyrate, propionate, and valerate, but lower molar fractions of others, including acetate and total branched VFAs (ANOVA, Table S3, Figure S3). Calves also tended to accumulate higher concentrations of fermentation intermediates like the VFA precursors ethanol, lactate, and succinate.

Overall, calves fed starter had the highest concentrations of total VFAs with both starter and mixed diet animals reaching total levels similar to adult cows (Figure S3). Concentrations and molar fractions of specific SCOAs varied across calf diet groups without clear trends consistent in any one diet. Specifically, silage-fed calves had the highest concentrations of formate and succinate, which was unlike adults, but had the lowest concentrations of ethanol, propionate, and valerate as well as the lowest molar proportions of propionate, which was similar to adults. Starter-fed calves were similarly inconsistent with high lactate concentrations common in calves but also high, adult-like acetate concentrations and valerate molar proportions (Figure S3). Calf butyrate concentrations occurred within the same range as those for adult cows with starter-fed calves having the highest concentrations among calves. However, butyrate molar proportions did not differ by any diet:age groups (Table S3).

Few specific OTUs correlated to SCOA concentrations and none correlated with molar fractions (Table S4). Strong correlations were mostly driven by age with calf associated OTUs being positively correlated with calf-associated fermentation intermediates. Specifically, calf-dominating *Methanobrevibacter* (A-OTU1) was positively associated with ethanol and lactate concentrations. Also, an unclassified fungal OTU found only in calves (F-OTU46) was positively associated with ethanol. Removing age as a factor to investigate possible dietary effects, only *Methanobrevibacter* (A-OTU1) remained strongly correlated with ethanol in calves alone (Table S4).

In contrast to the rumen microbiota, calf diet significantly correlated with overall fecal archaeal and bacterial communities in calves. At 4 weeks, corn silage and mixed diet archaeal communities differed in structure (PERMANOVA, Table S3). At weaning (8 weeks), archaeal and bacterial microbiota from corn silage-fed calves differed in both structure and composition from those fed starter or mixed diet (PERMANOVA, Table S3). Also, fecal archaeal communities of silage-fed calves had higher inter-animal variation at 8 weeks (PERMDISP, TukeyHSD, Table S3). Trends similar to those observed in rumen samples were observed for archaeal and fungal OTUs in feces, with more OTUs shared between adults and silage-fed calves compared to the other diets (Figure 3). Specific trends were not apparent for bacterial OTUs due to the high diversity and inter-animal variation among fecal samples at 8 weeks.

### Calf diet influences long-term rumen microbiota

Overall, all calves achieved an adult-like microbiota between weaning and 1-year of age, though additional differences were apparent between 1- and 2-year cows (Figure 1, Figure 2). Adult cows generally displaying lower beta-diversity than calves (Table S6), and calf diet significantly correlated with the structure and/or composition of some adult rumen but no fecal communities (PERMANOVA, Table S3). Overall, at 1 year, fungi differed in animals raised on corn silage and bacteria differed between all diets. At 2 years, archaea and bacteria differed in animals raised on calf starter while fungi differed between all diets (PERMANOVA, Table S3). Additionally, 2-year bacterial diversity was higher in the rumens of cows raised on silage relative to a mixed diet (ANOVA, TukeyHSD), and inter-animal variation in rumen bacterial, archaeal, and fungal communities varied by calf diet groups (PERMDISP, TukeyHSD, Table S3). No significant diet effects were observed in the structure, composition, variation, or diversity of communities in 1- or 2-year feces.

In general, the rumen communities of cows raised on different calf diets contained similar taxa but different abundant OTUs within these taxa. These included the bacterial taxa *Prevotella, Succiniclasticum*, and unclassified Succinivibrionaceae as well as the fungal taxa *Caeomyces, Piromyces*, and *Oripinomyces*; *Methanobrevibacter* archaeal OTUs also differed by diet groups. In contrast, silage-fed cows had more abundant *Fibrobacter* while starter-fed and mixed diet cows had highly abundant *Cyllamcyes*, both of which did not have abundant counterparts in the other diet groups.

Overall SCOA profiles and specific SCOAs were significantly different in 2-year-old cows raised on different diets (PERMANOVA, Table S3). Cows raised on the mixed diet had significantly different profiles (PERMANOVA) as well as higher concentrations of acetate and butyrate (ANOVA, Table S3, Figure S3). Also, branched VFAs isobutyrate and isovalerate + 2-methylbutyrate, which did not differ by diet in calves, were higher in mixed diet raised cows at 2 years. Molar fractions of VFAs did not differ between diet groups within adult animals.

### Calf diet not correlated with production measures

Animals gained weight from 2 to 8 weeks (*P* = 6.69E-30, 1.18 kg/day) and from 8 weeks to lactation (*P* = 1.35E-30, 0.456 kg/day) following linear trends with more accelerated growth in calves. Calf diet did not impact weight gain in calves (2 to 8 weeks, *P* = 0.432, Figure 4a) or in adults (8 weeks to > 2 years, *P* = 0.797, repeated measure linear regression, Figure 4b). Additionally, neither 2-year milk production efficiency (MPE, ANOVA *P* = 0.965) nor overall efficiency (*P* = 0.959) varied by calf diet (Figure 4c). No alpha- or beta-diversity measure of microbial communities significantly correlated with efficiency or MPE (PERMANOVA, ANOVA, Table S3). However, one unclassified Neocallimastigomycota (F-OTU19) displayed a strong positive correlation with efficiency (Kendall, Table S4). SCOA profile structure and composition correlated with MPE (PERMANOVA); however, no individual SCOAs significantly differed by milk production metrics (ANOVA, Table S3).

**Figure 4.**
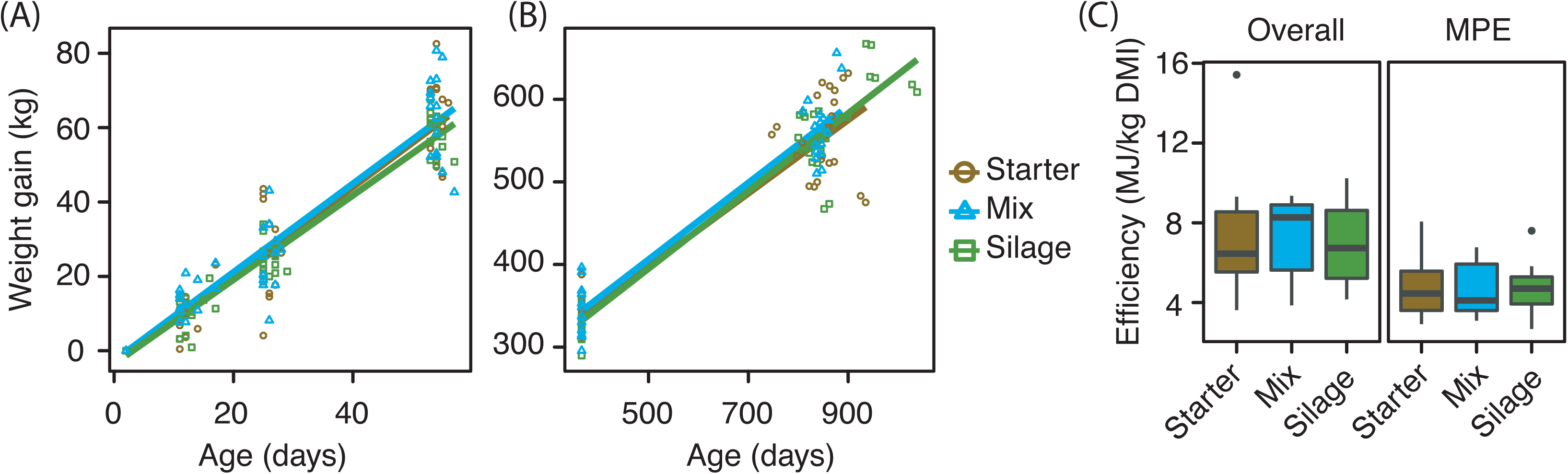
Calf diet impacts on weight gain and milk production. Weight gain (kg) of animals over time from (A) 2 days to weaning (8 weeks) and (B) 8 weeks to first lactation. Data were normalized to initial day 2 weights. (C) Milk and overall efficiency of dairy cows raised on different diets are also presented. Overall efficiency was determined by total energy expenditure (mastitis, gestation, weight maintenance and gain/loss, milk production, Table S8) per kg of dry matter intake (DMI). Milk production efficiency was determined by total milk production (MJ) per kg of DMI. Calf starter (brown), corn silage (green), mixture (blue).

### DISCUSSION

Purposeful alteration of GIT microbial communities has been proposed as a means to improve production of agriculturally relevant products like milk (Elie Jami *et al.*, 2014; Jewell *et al.*, 2015). Early-life interventions are a promising avenue as the developing microbiota in pre-weaned animals may be more easily altered than those in adults (Jami *et al.*, 2013; Yanez-Ruiz *et al.*, 2015; Dill-McFarland *et al.*, 2017). Here, we compared GIT microbiota and production traits of dairy cows raised on calf starter grains (investigated in depth previously (Dill-McFarland *et al.*, 2017)) to those raised on corn silage or a mixture of starter and silage in order to determine if early-life diet has long-term impacts on the adult cow.

Similar to calves raised on calf starter (Dill-McFarland *et al.*, 2017), GIT microbial communities in calves fed silage or mixed diets became more alike and more diverse with age. Animals had a succession of microorganisms representing similar taxa but different specific OTUs present at different ages. All animals achieved an adult-like microbiota between weaning (8 weeks) and one year of age. Therefore, age is a major driving force in the establishment of microbial communities in developing dairy calves, confirming previous reports (Jami *et al.*, 2013; Dill-McFarland *et al.*, 2017). However, in this study, diet-driven differences were also apparent, indicating that pre-weaning feed has a lesser, but still detectable, effect on the developing GIT microbiota.

While all calves progressed towards a similar adult microbiota, silage-fed animals achieved a more adult-like microbiota by weaning (8 weeks) compared to the other diets. Specifically, silage-fed calves at weaning had higher numbers and abundances of OTUs shared with adults, as well as fewer and less abundant OTUs strictly present in calves (Figure 3). At weaning, there was a striking difference in archaeal communities with silage-fed animals transitioning away from the calf-dominating *Methanobrevibacter* (A-OTU1) and lacking in all calf-associated unclassified fungi (Table S4).

Importantly, under typical modern production conditions, dairy calves receive minimal maternal care and have limited interaction with adult animals (Khan *et al.*, 2016). Calves are exposed to adult cow-associated microorganisms that are present in the environment or through contact with humans that interact with adult animals (Dill-McFarland *et al.*, 2017). Our data suggest that a calf diet of corn-silage may accelerate ruminal microbiota development into a more adult-like state by continually seeding the rumen with fiber, thus making complementary resources and nutrients readily available whenever exposure to adult-associated microbes occurs. In this way, corn-silage fed calves are likely able to maintain an adult-like ruminal microbiota well before full GIT development.

In addition, analysis of rumen short chain organic acids (SCOAs) indicated accelerated microbiota development in corn silage-fed calves at weaning. Specifically, silage was associated with lower concentrations of volatile fatty acid (VFA) intermediates, such as ethanol and lactate. These VFA intermediates generally do not accumulate in adult ruminants, as members of the microbiota convert them to VFAs readily absorbed by the host (Flint *et al.*, 2008). Conversions away from intermediates in silage-fed calves supports that a more complex, adult-like microbiota is present. Similar trends have been seen in calves reared with the addition of calf starter grains to a milk-only diet (Malmuthuge *et al.*, 2013), and the more pronounced results present in this study may be due to the silage’s greater similarity to adult feeds (Table S1), especially with respect to fiber content. Silage-fed calves also had low, adult-like concentrations and proportions of propionate, the main precursor of gluconeogenesis in ruminants (Zhang *et al.*, 2017). This is in contrast to starter and mixed diet calves, which had significantly higher propionate, indicative of reduced GIT development and a failure to meet nutritional needs from ingested feeds (Zhang *et al.*, 2017).

Some calf diet-driven differences in the microbiota were still present in adult animals, a phenomenon also observed in goats and lambs (Yáñez-Ruiz *et al.*, 2010; Abecia *et al.*, 2013, 2014). Similar to age-related changes (Dill-McFarland *et al.*, 2017), overall community structures (Bray-Curtis) differed and comparable taxa but different OTUs were present in adult cows raised on different diets. The presence or abundance of adult-associated OTUs at weaning had no correlation to actual adult OTU abundances, and no significant differences in OTU abundances in calves persisted in the same OTUs in adults. Therefore, the apparent long-term effects of calf diet on adult microbial communities appear to be weak and indirect. These effects may be mediated by differences in internal development, as supplements have been shown to impact rumen size, weight, and/or papillation (Warner *et al.*, 1956; Tamate *et al.*, 1962; Suárez *et al.*, 2007; Suarez-Mena *et al.*, 2011) or by differential programming of the immune system by early-life microorganisms (Hooper *et al.*, 2012; Gensollen *et al.*, 2016). Taken together, this indicates that early-life feeds aimed at physiological and/or immune system develop may be more impactful on later adult microbiota than those specifically targeted at microorganisms.

Although calf diet correlated with the rumen microbiota of adult cows, it did not significantly impact weight gain, milk production, or overall efficiency. Additionally, no adult microbial communities differed by calf diet. This indicates that the weaning transition, and factors after weaning, significantly contribute to the establishment of an adult-like microbiota and may allow under-developed animals to “catch up” such that early-life differences are no longer apparent or impactful on production. Thus, we propose that efforts to directionally alter the microbiota toward improved production and efficiency should be explored in calves during the weaning transition, prior to the stabilization of host-specific communities observed in adult cows.

Overall, this work concurrently evaluates the effects of pre-weaning diet on growth, milk production, and associated GIT microbial communities in dairy cows over time. While it does not appear that pre-weaning diet can be used to promote specific microorganisms or improve efficiency in the adult cow, there were no significant detriments to feeding calves corn silage over commercially prepared calf starter grains. Given that corn silage is often significantly cheaper than calf starter grains and more readily accessible on dairy farms, feeding corn silage to calves could result in substantial cost savings for dairy producers. Our study also supports that pre-weaning calf management appears to be most effective at promoting physiological development and animal growth, as has historically been the case, while the weaning transition appears to be an opportune time for altering microbial communities for long-term production gains. These results have implications for management and feeding practices across numerous agricultural systems as more efficient, less environmentally costly food production is needed to meet global demands.

## EXPERIMENTAL PROCEDURES

### Animals and diet

This study was completed under animal use protocol A01501, approved by the College of Agriculture and Life Sciences’ Institutional Animal Care and Use Committee, University of Wisconsin-Madison. Three cohorts of 15 (3 males, 12 females) Holstein dairy calves each (total N = 45) were raised at the US Dairy Forage Research Center (USDFRC) Research Farm near Prairie du Sac, WI. All calves received pasteurized milk with milk balancer protein-blend (Land O’Lakes, St. Paul, MN) added to 15% total milk solids. From 0 to 5 days of age, animals were fed 6 quarts (∼5.7 L) of milk per day and from 6 days until weaning (approximately 56 days), 8 quarts (∼7.6 L) per day. Each cohort was also offered *ad libitum* access to one of three randomly assigned supplemental feeds: calf starter (diet A; 58.25% whole corn, 1.75% molasses, and 40% Future Cow Ampli-Calf Mixer Pellet B150, Purina Animal Nutrition, Shoreview, MN) (Dill-McFarland *et al.*, 2017), corn silage (diet B; USDFRC), or a 25:75 by weight mixture of calf starter and corn silage respectively (diet C; approximately 50:50 by dry matter). Detailed nutritional analyses of feeds are provided (Table S1). Supplement offered to and refused by each calf was weighed daily to determine supplement intake (Table S7).

After weaning, calves were transitioned through a series of standard heifer diets until placed on the lactating herd total mixed ration (TMR) diet after calving (Table S1). TMR offered to and refused by each cow was weighed and subsamples were taken daily during the milk sampling period (150 to 158 days in milk [DIM]) to determine nutrient intake using established methods (Weimer *et al.*, 1999). All animals had *ad libitum* access to water throughout the trial.

### Rumen sampling and feces collection

Samples were collected between June 2012 and July 2015. The 9 male calves (3 per diet cohort) were sacrificed near weaning (56 – 68 days) to obtain rumen samples. Rumen liquids were obtained by straining total rumen contents through four layers of cheesecloth. Solids remaining in the cheesecloth were squeezed to remove all free liquid and transferred to a separate sterile container. Twelve heifers were ruminally cannulated (9.5 – 11 months, 4 per diet cohort, animal use protocol A01307), and rumen liquids and solids were collected through the cannula before morning feeding on three consecutive days at 1 year (365 ± 7 days) and in the middle of the first lactation cycle after two years (154 – 156 DIM). All samples were immediately transported on wet ice and stored at -80 °C prior to DNA extraction.

Fresh feces were obtained by hand from the rectum of animals using clean nitrile gloves. Samples were taken at 2 weeks (14 ± 3 days, N = 40) or 3 weeks (21 ± 2 days, N = 5) in an effort to avoid sampling during illness. Feces were also collected from all animals at 4 weeks (27 ± 2 days), 8 weeks (54 ± 2 days), 1 year (365 ± 7 days), and the middle of first lactation (155 ± 1 DIM). Samples were stored at -20 °C on site, transported on wet ice, and then stored at -80 °C prior to DNA extraction.

### Milk collection

Total milk production was measured for each cow over 9 consecutive days during the middle of the first lactation cycle (151 – 159 DIM) offset one day from feed sampling as morning milk production is impacted by the previous day’s feed. Milk samples were collected from all three daily milkings on these days. Milk was stored at 4 °C and submitted to AgSource Cooperative Services (Verona, WI) for near-infrared spectroscopic prediction to determine composition including fat, lactose, protein, non-fat solids, milk urea nitrogen, and somatic cell count (Tsenkova *et al.*, 1999).

### Growth and health

Cow health was assessed by standard daily monitoring for disease including fecal and attitude scores as described (Dill-McFarland *et al.*, 2017). Calves were treated with antibiotics and electrolytes for scours (diarrhea) and with antibiotics plus fever reducer for respiratory disease, as required. Cows were treated for ketosis with propylene glycol to restore energy balance and for mastitis with antibiotics, as necessary. Calf fecal samples taken during treatment for scours (N = 16) or respiratory disease (N = 3) are noted (Table S2). No samples were taken during illness among adult animals.

Calves were weighed at 2 days and near the time of fecal sampling from 2 to 8 weeks. Cows were weighed on three consecutive days at 1 year (365 – 367 days) and 2 sets of 3 days bracketing the milk sampling period during first lactation (149 – 151 and 159 – 161 DIM).

### SCOA analysis

Analysis of short-chain organic acids (SCOAs) was performed on rumen liquids following standard methods using high-performance liquid chromatography (HPLC) (Weimer *et al.*, 1991). In brief, a portion of rumen liquid samples were treated with calcium hydroxide and cupric sulfate and then run on an Aminex HPX-87H HPLC column (Bio-Rad, Hercules, CA, USA) with a Pro-Star autosampler (Varian, Palo Alto, CA, USA) and refractive index detector (Knauer, Berlin, Germany). The flow rate of the 0.015 N H2SO4/0.0034 M EDTA mobile phase was 0.70 ml/min at 45 °C. Samples were compared to two standard curves: one containing 10 mM concentrations of acetate, butyrate, formate, isobutyrate, isovalerate, 2-methylbutyrate, propionate, succinate and valerate, and one containing 10 mM lactate. Standards were run for every 10 samples, and crotonate (2-butenoate) was added to all samples as an internal standard. Separate analyses confirmed that detector response for each individual SCOA was linear to at least 300 mM.

Total volatile fatty acid (VFA) concentrations were calculated as the sum of acetate, propionate, isobutyrate, butyrate, isovalerate, 2-methylbutyrate, and valerate. Total branched chain VFAs included isobutyrate, isovalerate and 2-methylbutyrate. Molar fractions (mF) of VFAs were also calculated compared to total VFA concentrations per sample.

### DNA extraction and sequencing

Total genomic DNA was extracted from samples following a mechanical disruption and hot/cold phenol extraction protocol published previously (Stevenson and Weimer, 2007) with feces processed in a manner similar to rumen solids and with the following modification: 25:24:1 phenol:chloroform:isoamyl alcohol was used in place of phenol:chloroform and samples required up to six additional washes. All DNA samples were resuspended in water, quantified by a Qubit® Fluorometer (Invitrogen, San Diego, CA, USA), and stored at -20 °C.

For bacteria, PCR was performed using universal primers flanking the variable 4 (V4) region of the bacterial 16S rRNA (Kozich *et al.*, 2013). Conditions, purification, and sequencing were as previously described (Dill-McFarland *et al.*, 2017). Briefly, PCR was performed using 50 ng of DNA with HotStart ReadyMix (KAPA Biosystems, Wilmington, MA, USA) and products were purified by gel extraction (Zymo Research, Irvine, CA). Samples were quantified by Qubit® Fluorometer, equimolar pooled, and sequenced with the MiSeq 2×250 v2 kit (Illumina, San Diego, CA, USA) using custom sequencing primers (Kozich *et al.*, 2013).

For archaea and fungi, a two-step PCR protocol was employed as previously described (Dill-McFarland *et al.*, 2017). In short, the archaeal V6-8 16S region or the fungal internal transcribed region 1 (ITS1) was amplified (Kittelmann *et al.*, 2013) and Illumina adapters with unique indices were added. PCR products were column-purified (PureLink, Invitrogen) after the first PCR and gel-extracted (Zymo Research) after the second. Samples were quantified by Qubit® Fluorometer, equimolar pooled, and sequenced with the MiSeq 2×300 v3 kit (Illumina). All DNA sequences were deposited in the NCBI’s Sequence Read Archive under the accession no. PRJNA319127.

### Sequence clean-up

All sequences were demultiplexed on the Illumina MiSeq. Further sequence processing was performed using mothur v.1.36.1 (Schloss *et al.*, 2009) following a protocol adapted from (Kozich *et al.*, 2013). The full clean-up pipeline is detailed in Supplementary Text S1 in (Dill-McFarland *et al.*, 2017). Briefly, paired-end sequences were combined into contigs and poor-quality sequences were removed. Bacterial and archaeal sequences were aligned against the SILVA 16S rRNA gene reference alignment database (Pruesse *et al.*, 2007). Fungal sequences were aligned *de novo* within the dataset. For all amplicons, sequences were pre-clustered and chimera detection and removal was performed. Bacterial and archaeal sequences were classified to the GreenGenes database (DeSantis *et al.*, 2006); fungal sequences were classified to the UNITE dynamic ITS database (Kõljalg *et al.*, 2013). Singletons were removed to facilitate downstream analyses.

All sequences were grouped into 97% operational taxonomic units (OTUs) by uncorrected pairwise distances and furthest neighbor clustering. Coverage was assessed by Good’s coverage calculated in mothur. Bacterial (9 000 seqs), archaeal (100), and fungal (250) communities were normalized to equal sequence counts near their lowest respective sample, and these normalized OTU tables were used in all further analyses.

### Animal statistics

All tests were assessed at a significance of *P* < 0.05. Differences between diet groups were assessed by repeated measures linear regression over time and by calf for weight gain and supplement intake with the lme4 (Bates *et al.*, 2015) and lmerTest (Kuznetsova *et al.*, 2016) packages in R v3.2.3 (R Core Team, 2017). Weight gain was normalized to day 2 weights, and supplement intake was log-transformed. Milk production efficiency was calculated based on energy-corrected milk (ECM) production, body weight change and maintenance, gestation, and mastitis infection as described (Table S8) (Council, 2001; Dairy Records Management Systems, 2013). ECM from the three daily milkings were summed per day, and dry matter intake (DMI) was calculated as feed offered minus feed refused in kg. Animal weights before and after the milk sampling period were averaged across three days each. Daily ECM, DMI, and milk somatic cell count (SCC) were averaged across the period for each cow. The energetic cost of mastitis was determined based on SCC in milk converted to potential milk production loss (Dairy Records Management Systems, 2013). Efficiency was defined as total energy output (MJ) of the five energetic demands (*i.e.* ECM, weight maintenance, weight change, gestation, mastitis) per kg of DMI. Efficiencies were compared between diet groups with ANOVA and correlated to OTUs at a minimum of 0.5% relative abundance in at least one sample using Kendall’s correlations with microbial data averaged across consecutive sampling days to avoid animal effects. All code is available at https://github.com/kdillmcfarland/GS01.

### Microbiota statistics

Beta-diversity was visualized by nonmetric multidimensional scaling (nMDS) plots of the Bray-Curtis metric calculated with square root transformed data in R (vegan package (Oksanen *et al.*, 2015)). Differences in the spread of Bray-Curtis values within age and diet groups were calculated by permutation tests of multivariate homogeneity of group dispersions (PERMDISP, vegan) and pairwise between groups with Tukey’s HSD.

All tests were assessed at a significance of *P* < 0.05, and strong correlations were defined as either > 0.7 or < -0.7. Alpha-diversity was determined with Shannon’s diversity and Chao’s richness calculated in mothur. Differences in community diversity and richness were assessed overall by ANOVA and pairwise between groups within significant ANOVAs by Tukey’s HSD from multiple comparisons in R. All samples were modeled for calf diet, age group, and diet within age groups (diet:age group) (Model 1). Two-year samples were modeled for milk and overall efficiency (Model 2) with rumen OTU tables averaged across the 3 consecutive days at 1 or 2 years to avoid animal effects.

Total structure (relative abundance, Bray-Curtis) and composition (presence/absence, Jaccard) of OTUs and SCOA concentrations were evaluated for changes using permutational ANOVA (PERMANOVA, vegan) with the same models as ANOVA (Model 1 and 2). Also, fecal OTUs at 2- and 4-weeks were modeled by scours versus healthy since some of these samples were taken during illness (Model 3). Pairwise comparisons between groups within significant PERMANOVAs were completed using PERMANOVAs with a Benjamini-Hochberg correction for multiple comparisons. OTUs contributing to differences between groups in PERMANOVA were identified by analysis of similarity percentages (SIMPER, vegan). OTUs contributing at least 1% of the SIMPER variation were assessed further using Kruskal-Wallis with the Benjamini-Hochberg correction for multiple comparisons.

Co-variation of the microbiota with SCOAs in rumen liquids was tested using Mantel tests for dissimilarity matrices. Differences between SCOA concentrations and molar fractions within age groups and diets were assessed overall by ANOVAs with a Benjamini-Hochberg correction for multiple comparisons and pairwise between groups with TukeyHSD. OTUs with at least 0.5% relative abundance in one or more samples were associated with SCOAs by Kendall’s correlations to 3-day averaged rumen OTU tables. All code is available at https://github.com/kdillmcfarland/GS01.

## ACKNOWLEDGEMENTS

The authors wish to thank the staff at the USDFRC for daily animal care, and in particular, Ron Skoyen for his assistance. The authors also thank all members of the Suen laboratory for their support and careful reading of the manuscript. Finally, we would like to thank the Wisconsin Energy Institute’s IT group for access to computational resources. This work was supported by U.S. Department of Agriculture National Institute of Food and Agriculture HATCH grant WIS01729 and foundational grant 2015-67015-23246 to GS, and a USDA Agricultural Research Service CRIS project 5090-21000-024-00D to PJW. The authors declare no conflict of interest.

## FIGURE LEGENDS

**Figure S1. Calf supplement intake.** Supplement intake (g/day) by diet from birth to weaning. A) Raw supplement intake. B) Log-transformed supplement intake to fit a linear model. Calf starter (brown, circle), corn silage (green, square), mixture (blue, triangle).

**Figure S2. Richness of the microbiota in cows raised on different diets.** Chao’s richness of archaeal, bacterial, and fungal communities in feces as well as rumen liquids and solids. Boxes are colored by animal age for 2-week (red), 4-week (orange), 8-week (yellow), 1-year (green), and 2-year (blue). Ages with significantly different richness across each amplicon are indicated by different letters. Asterisks denote groups containing significant diet differences (TukeyHSD, *P* < 0.05, Table S3).

**Figure S3. Short chain organic acid (SCOA) profiles.** (A) SCOA concentrations and (B) VFA molar fractions in rumen liquids at 8 weeks, 1 year, and 2 years. Boxes are colored by animal age for 8-weeks (yellow), 1-year (green), and 2-years (blue). Letters within each plot denote significantly different groups (TukeyHSD, Table S3). Calf starter (St), corn silage (Si), mixed diet (M).

## REFERENCES

Abecia, L., Martín-García, A.I., Martínez, G., Newbold, C.J., and Yáñez-Ruiz, D.R. (2013) Nutritional intervention in early life to manipulate rumen microbial colonization and methane output by kid goats postweaning. J. Anim. Sci. 91:.

Abecia, L., Waddams, K.E., Martínez-Fernandez, G., Martín-García, A.I., Ramos-Morales, E., Newbold, C.J., and Yáñez-Ruiz, D.R. (2014) An antimethanogenic nutritional intervention in early life of ruminants modifies ruminal colonization by archaea. Archaea 2014: 841463.

Bajagai, Y.S., Klieve, A. V., Dart, P.J., and Bryden, W.L. (2016) Probiotics in animal nutrition - Production, impact and regulation No. 179. Makkar, H.P.S. (ed) Food and Agricultural Organization of the United Nations, Rome.

Bates, D., Maechler, M., and Bolker, B. (2015) Fitting linear mixed-effects models using lme4. J. Stat. Softw. 67: 1–48.

Bergman, E.N. (1990) Energy contributions of volatile fatty acids from the gastrointestinal tract in various species. Physiol. Rev. 70: 567–590.

Cannon, S.J., Fahey Jr, G.C., Pope, L.L., Bauer, L.L., Wallace, R.L., Miller, B.L., and Drackley, J.K. (2010) Inclusion of psyllium in milk replacer for neonatal calves. 2. Effects on volatile fatty acid concentrations, microbial populations, and gastrointestinal tract size. J. Dairy Sci. 93: 4744–4758.

Carberry, C.A., Kenny, D.A., Han, S., McCabe, M.S., and Waters, S.M. (2012) Effect of phenotypic residual feed intake and dietary forage content on the rumen microbial community of beef cattle. Appl. Environ. Microbiol. 78: 4949–4958.

Council, N.R. ed. (2001) Nutrient requirements of dairy cattle. Dairy Records Management Systems (2013) The DHI Glossary Raleigh, NC.

DeSantis, T.Z., Hugenholtz, P., Larsen, N., Rojas, M., Brodie, E.L., Keller, K., et al. (2006) Greengenes, a chimera-checked 16S rRNA gene database and workbench compatible with ARB. Appl. Environ. Microbiol. 72: 5069–5072.

Dill-McFarland, K.A., Breaker, J.D., and Suen, G. (2017) Microbial succession in the gastrointestinal tract of dairy cows from 2 weeks to first lactation. Sci. Rep. 7: 40864.

Flint, H.J., Bayer, E.A., Rincon, M.T., Lamed, R., and White, B.A. (2008) Polysaccharide utilization by gut bacteria: potential for new insights from genomic analysis. Nat Rev Micro 6: 121–131.

Fonty, G., Gouet, P., Jouany, J.-P., and Senaud, J. (1987) Establishment of the microflora and anaerobic fungi in the rumen of lambs. Microbiology 133: 1835–1843.

Gensollen, T., Iyer, S.S., Kasper, D.L., and Blumberg, R.S. (2016) How colonization by microbiota in early life shapes the immune system. Science 352: 539–544.

Hooper, L. V, Littman, D.R., and Macpherson, A.J. (2012) Interactions between the microbiota and the immune system. Science 336: 1268–1273.

Jami, E., Israel, A., Kotser, A., and Mizrahi, I. (2013) Exploring the bovine rumen bacterial community from birth to adulthood. ISME J. 7: 1069–1079.

Jami, E., Shterzer, N., Yosef, E., Nikbachat, M., Miron, J., and Mizrahi, I. (2014) Effects of including NaOH-treated corn straw as a substitute for wheat hay in the ration of lactating cows on performance, digestibility, and rumen microbial profile. J. Dairy Sci. 97: 1623–1633.

Jami, E., White, B.A., and Mizrahi, I. (2014) Potential role of the bovine rumen microbiome in modulating milk composition and feed efficiency. PLoS One 9: e85423.

Jatkauskas, J. and Vrotniakiene, V. (2010) Effects of probiotic dietary supplementation on diarrhoea patterns, faecal microbiota and performance of early weaned calves. Vet. Med. (Praha). 55: 494–503.

Jewell, K.A., McCormick, C.A., Odt, C.L., Weimer, P.J., and Suen, G. (2015) Ruminal bacterial community composition in dairy cows is dynamic over the course of two lactations and correlates with feed efficiency. Appl. Environ. Microbiol. 81: 4697–4710.

Khan, M.A., Bach, A., Weary, D.M., and von Keyserlingk, M.A.G. (2016) Invited review: Transitioning from milk to solid feed in dairy heifers. J. Dairy Sci. 99: 885–902.

Kittelmann, S., Seedorf, H., Walters, W.A., Clemente, J.C., Knight, R., Gordon, J.I., and Janssen, P.H. (2013) Simultaneous amplicon sequencing to explore co-occurrence patterns of bacterial, archaeal and eukaryotic microorganisms in rumen microbial communities. PLoS One 8: e47879.

Klein-Jöbstl, D., Schornsteiner, E., Mann, E., Wagner, M., Drillich, M., and Schmitz-Esser, S. (2014) Pyrosequencing reveals diverse fecal microbiota in Simmental calves during early development. Front. Microbiol. 5: 622.

Kõljalg, U., Nilsson, R.H., Abarenkov, K., Tedersoo, L., Taylor, A.F.S., Bahram, M., et al. (2013) Towards a unified paradigm for sequence-based identification of fungi. Mol. Ecol. 22: 5271–5277.

Kozich, J.J., Westcott, S.L., Baxter, N.T., Highlander, S.K., and Schloss, P.D. (2013) Development of a dual-index sequencing strategy and curation pipeline for analyzing amplicon sequence data on the MiSeq Illumina sequencing platform. Appl. Environ. Microbiol.

Kuznetsova, A., Brockhoff, P.B., and Christensen, R.H.B. (2016) Tests in linear mixed effects models (lmerTest).

Lallès, J.P. (2012) Long term effects of pre- and early postnatal nutrition and environment on the gut1. J. Anim. Sci. 90: 421–429.

Malmuthuge, N., Griebel, P.J., and Guan, L.L. (2015) The gut microbiome and its potential role in the development and function of newborn calf gastrointestinal tract. Front. Vet. Sci. 2:.

Malmuthuge, N., Li, M., Goonewardene, L.A., Oba, M., and Guan, L.L. (2013) Effect of calf starter feeding on gut microbial diversity and expression of genes involved in host immune responses and tight junctions in dairy calves during weaning transition. J. Dairy Sci. 96: 3189–3200.

McFall-Ngai, M., Hadfield, M.G., Bosch, T.C.G., Carey, H. V, Domazet-Lošo, T., Douglas, A.E., et al. (2013) Animals in a bacterial world, a new imperative for the life sciences. Proc. Natl. Acad. Sci. U. S. A. 110: 3229–3236.

de Menezes, A.B., Lewis, E., O’Donovan, M., O’Neill, B.F., Clipson, N., and Doyle, E.M. (2011) Microbiome analysis of dairy cows fed pasture or total mixed ration diets. FEMS Microbiol. Ecol. 78: 256–265.

Oksanen, J., Blanchet, F.G., Kindt, R., Legendre, P., Minchin, P.R., O’Hara, R.B., et al. (2015) vegan: community ecology package.

Patra, A.K. and Yu, Z. (2012) Effects of essential oils on methane production and fermentation by, and abundance and diversity of, rumen microbial populations. Appl. Environ. Microbiol. 78: 4271–4280.

Pruesse, E., Quast, C., Knittel, K., Fuchs, B.M., Ludwig, W., Peplies, J., and Glockner, F.O. (2007) SILVA: a comprehensive online resource for quality checked and aligned ribosomal RNA sequence data compatible with ARB. Nucleic Acids Res. 35: 7188–7196.

R Core Team (2017) R: A language and environment for statistical computing.

Schloss, P.D., Westcott, S.L., Ryabin, T., Hall, J.R., Hartmann, M., Hollister, E.B., et al. (2009) Introducing mothur: open-source, platform-Independent, community-supported software for describing and comparing microbial communities. Appl. Environ. Microbiol. 75: 7537–7541.

Signorini, M.L., Soto, L.P., Zbrun, M. V, Sequeira, G.J., Rosmini, M.R., and Frizzo, L.S. (2012) Impact of probiotic administration on the health and fecal microbiota of young calves: A meta-analysis of randomized controlled trials of lactic acid bacteria. Res. Vet. Sci. 93: 250–258.

Soberon, F., Raffrenato, E., Everett, R.W., and Van Amburgh, M.E. (2012) Preweaning milk replacer intake and effects on long-term productivity of dairy calves. J. Dairy Sci. 95: 783–793.

Stevenson, D. and Weimer, P. (2007) Dominance of Prevotella and low abundance of classical ruminal bacterial species in the bovine rumen revealed by relative quantification real-time PCR. Appl. Microbiol. Biotechnol. 75: 165–174.

Suarez-Mena, F.X., Hill, T.M., Heinrichs, A.J., Bateman, H.G., Aldrich, J.M., and Schlotterbeck, R.L. (2011) Effects of including corn distillers dried grains with solubles in dairy calf feeds. J. Dairy Sci. 94: 3037–3044.

Suárez, B.J., Van Reenen, C.G., Stockhofe, N., Dijkstra, J., and Gerrits, W.J.J. (2007) Effect of roughage source and roughage to concentrate ratio on animal performance and rumen development in veal calves. J. Dairy Sci. 90: 2390–2403.

Tamate, H., McGilliard, A.D., Jacobson, N.L., and Getty, R. (1962) Effect of various dietaries on the anatomical development of the stomach in the calf. J. Dairy Sci. 45: 408–420.

Torok, V.A., Percy, N.J., Moate, P.J., and Ophel-Keller, K. (2014) Influence of dietary docosahexaenoic acid supplementation on the overall rumen microbiota of dairy cows and linkages with production parameters. Can. J. Microbiol. 60: 267–275.

Tsenkova, R., Atanassova, S., Toyoda, K., Ozaki, Y., Itoh, K., and Fearn, T. (1999) Nearinfrared spectroscopy for dairy management: Measurement of unhomogenized milk composition. J. Dairy Sci. 82: 2344–2351.

Vlková, E., Rada, V., Trojanová, I., Killer, J., Šmehilová, M., and Molatová, Z. (2008) Occurrence of Bifidobacteria in faeces of calves fed milk or a combined diet. Arch. Anim. Nutr. 62: 359–365.

Wallace, R.J., Rooke, J.A., McKain, N., Duthie, C.-A., Hyslop, J.J., Ross, D.W., et al. (2015) The rumen microbial metagenome associated with high methane production in cattle. BMC Genomics 16: 839.

Warner, R.G., Flatt, W.P., and Loosli, J.K. (1956) Dietary factors influencing the development of the ruminant stomach. J. Agric. Food Chem. 4: 788–792.

Weimer, P., Shi, Y., and Odt, C. (1991) A segmented gas/liquid delivery system for continuous culture of microorganisms on insoluble substrates and its use for growth of Ruminococcus flavefaciens on cellulose. Appl. Microbiol. Biotechnol. 36: 178–183.

Weimer, P.J., Stevenson, D.M., Mantovani, H.C., and Man, S.L.C. (2010) Host specificity of the ruminal bacterial community in the dairy cow following near-total exchange of ruminal contents. J. Dairy Sci. 93: 5902–5912.

Weimer, P.J., Waghorn, G.C., Odt, C.L., and Mertens, D.R. (1999) Effect of diet on populations of three species of ruminal cellulolytic bacteria in lactating dairy cows. J. Dairy Sci. 82: 122–134.

Yanez-Ruiz, D.R., Abecia, L., and Newbold, C.J. (2015) Manipulating rumen microbiome and fermentation through interventions during early life: a review. Front. Microbiol. 6:.

Yáñez-Ruiz, D.R., Macías, B., Pinloche, E., and Newbold, C.J. (2010) The persistence of bacterial and methanogenic archaeal communities residing in the rumen of young lambs. FEMS Microbiol. Ecol. 72: 272–278.

Zhang, Q., Koser, S.L., Bequette, B.J., and Donkin, S.S. (2017) Effect of propionate on mRNA expression of key genes for gluconeogenesis in liver of dairy cattle. J. Dairy Sci. 98: 8698–8709.

